# Evidence for Philopatry and Homing in a New World Elapid Snake

**DOI:** 10.1101/092833

**Authors:** Alexander S. Hall, Abigail M. K. Vázquez-Quinto, Antonio Ramírez-Velázquez, Eric N. Smith

**Affiliations:** Department of Biology, The University of Texas at Arlington, Arlington, TX 76019, USA.; Zoológico Regional Miguel Álvarez del Toro, Calzada Cerro Hueco, Col Zapotal, C. P. 29094, Tuxtla Gutiérrez, Chiapas, México.

**Keywords:** compass sense, *Micrurus apiatus*, range maintenance, snake, coralsnake, Elapidae, Chiapas, philopatry, homing

## Abstract

Animal navigation allows individuals to efficiently find and use best available habitats. Despite the long history of research into well-studied taxa (e.g., pigeons, salmon, sea turtles), we know relatively little about squamate navigational abilities. Among snakes, documented philopatry (range maintenance) in a non-colubrid species has been rare. In this study, we document the first example of philopatry and homing in a new world elapid snake, *Micrurus apiatus*. Our data come from the first multi-year mark-recapture study of this species at the open urban preserve *Zoológico Regional Miguel Álvarez del Toro*, in Tuxtla Gutiérrez, Chiapas, Mexico. We show that on average snakes returned to within 144 m of their last capture point. By releasing snakes in one location, we noted that recaptured individuals preferentially returned to their last capture location, compared to a distribution of random locations in the park. We conclude with a preliminary discussion of the evolution of snake homing and potential mechanisms.

## Introduction

An animal’s use of available information to willfully move within its environment can be considered navigation (Lockley, 1967). Navigation, in the broadest sense, allows animals to efficiently utilize the resources available to them. The best descriptions of animal vertebrate navigation come from a handful of species: homing pigeons (e.g., Mora et al., 2004), salmon (reviewed in Dittman & Quinn, 1996), and sea turtles (e.g., Lohmann et al., 2004; Shimada et al., 2016). Each of these species spectacularly migrates hundreds, sometimes thousands, of kilometers from known ‘home’ locations and returns to these location with exceptional accuracy. Ethologists use focal species to describe the nature of navigation (Tinbergen, 1984: 58–79), but focusing on these species may ignore additional diversity of sensory mechanisms and behaviors available for navigation to other animals.

Among vertebrates, navigational ability in amphibians and non-avian reptiles remains understudied. This may be due to the relatively smaller size of most species and difficulty with detection when compared to more visible vertebrates. These animals rely on moving in and about narrowly defined microhabitats for shelter, feeding, reproduction, etc.; thus, one expects spatial navigation to provide fitness-associated benefits. Indeed, displaced newts *(Taricha rivularis*) travel over 8 km to their home streams using olfactory cues (Twitty, Grant, & Anderson, 1964; Grant, Anderson, & Twitty, 1968). Similarly, red spotted newts *(Notophthalmus viridescens)* orient towards their home location when displaced by about 45 km (Fischer et al., 2001). Even displaced frogs that do not require bodies of water to breed *(Eleutherodactylus coqui*) return to home territories (e.g., Gonser & Woolbright, 1995). Amphibians generally require damp microhabitats to avoid desiccation and are thus strongly reinforced for finding suitable habitats. Reptiles, on the other hand, can generally avoid water sources for extended periods.

Among squamates (lizards and snakes), we know even less about navigational capabilities and homing behavior. Several studies detail movement and migration in snakes, but few comment on actual navigational capabilities. Ciofi and Chelazzi (1994) measured homing patterns in a whip snake (*Hierophis viridiflavus*) and showed these snakes home to a primary shelter (i.e., philopatry) after excursions lasting a day to a month. From observation of a single animal, Singh et al. (1992) suggested homing behavior in Burmese pythons *(Python molurus)*. A more recent investigation of homing capability in *P. molurus*, populations introduced to the Florida everglades, determined that individuals displaced several (>30) kilometers from their initial capture location returned to within 5 km, with consistent movement across several weeks (Pittman et al., 2014). To date, all but one known occurrences of homing in terrestrial snakes have been described from the family Colubridae (Ciofi and Chelazzi, 1994) – two-thirds of all snake species (Uetz, 2016).

A homing study by Shetty and Shine (2002) on the yellow-lipped sea kraits *(Laticauda colubrina)* demonstrated that these elapids (i.e., non-colubrid) move to forage between two Fiji islands 5.3 km apart; however they always return to their home island. The philopatric, or range maintenance, behavior was observed within the span of several weeks. The kraits also exhibited philopatry between years and in juvenile snakes (Shetty & Shine, 2002). These authors also refer to their unpublished work on *L. colubrina* and another elapid sea snake, *L. frontalis*, where four of 26 juveniles returned to a capture site 5 and 12 days after being translocated 4 km. Though a handful of studies on terrestrial elapids report home range sizes and preferred microhabitats (Angilletta Jr, 1994: green mamba, *Dendroapsis angusticeps;* Akani et al., 2005: Goldie’s tree cobra, *Pseudohaje goldii;* and Croak et al., 2013: Australian broad-headed snake, *Hoplocephalus bungaroides)*, we are aware of only one suggesting homing behavior (Rao et al., 2012: based on a single king cobra, *Ophiophagus hannah)*, but none demonstrating an elapid repeatedly returning home after being translocated. In the Agumbe reserve in India, a male translocated king cobra was radio-tracked for more than 100 km and then lost, apparently due to homing (Bhaisare et al. 2010; Rao et al. 2012, 2013). In this study, we performed a mark-recapture study of an elapid snake, *Micrurus apiatus* (formerly *Micrurus browni browni*; Schmidt & Smith, 1943, Reyes-Velasco, 2015), and discovered another instance of homing behavior in the family Elapidae: one from a fully terrestrial species and the first example from a new world elapid.

The venomous monadal coralsnake population from the valley of Tuxtla Gutiérrez, Chiapas, has traditionally been considered the nominal subspecies of *M. browni* (Campbell & Lamar, 2004: 153-155), but it was recently shown to be unrelated to this species and should be better known as *M. apiatus* (Reyes-Velasco, 2015). Reyes-Velasco (2015) based this determination on the combined analysis of mitochondrial data and over 1,000 independent nuclear loci. These data contradicts traditional taxonomy differentiating coralsnakes in the region based on the presence or absence of supracloacal keels. It appears that supracloacal keels are usually present in arid or seasonal regions. Based on our knowledge of the species as *M. b. browni, M. apiatus* is found in moderately high abundance in southern Mexico and southwestern Guatemala (Álvarez del Toro, 1982; Campbell & Lamar, 1989; Köhler, 2008; Reyes-Velasco, 2015). Cryptic behavior (thus low detectability) keeps the ecology and behavior of most *Micrurus* species poorly understood (Campbell & Lamar, 2004), with only a few published reports on *Micrurus* ecology and behavior (e.g., Greene, 1984; Marques & Sazima, 1997).

Via daily opportunistic sampling in an urban open park, we capitalized on a unique opportunity to conduct the first mark-recapture study of *M. apiatus*. From our records, we report on sex-specific movement patterns of this population of *M. apiatus*. Captured snakes released at a standard location just outside the park sometimes revisited their original capture location. We thus report on homing behavior by these snakes.

## Methods

We collected *Micrurus apiatus* within the Miguel Álvarez del Toro Zoo (ZooMAT) – an open zoological preserve available daily to the public and located on a hillside in tropical Chiapas, Mexico, 2 km southwest of the city of Tuxtla Gutiérrez (16°43’N, 93°06’W). Park support staff walked the 2.5 km of trails daily, in the morning and evening between 0700 and 1800 hours. These staff reported *M. apiatus* sightings to the authors (AMKVQ and ARV), who transported the animal in a polyethylene bag to the reptile and amphibian building for examination and marking. When reporting sightings, park staff recorded the date, time, and exact location of the encounter. For initial encounters with each snake, we recorded: cloacal temperature (°C), total body length (mm), tail length (mm), number of red rings, number of caudal yellow rings, mass (g), sex, and ambient outdoor temperature (°C). After examination we tattooed, using a sterile syringe and black India ink, injecting under posterior red ventral scales. To reduce the amount of ink applied per animal, we used separate numbering for males and females, so that low value numbers could be reused.

Because we found *M. apiatus* in public areas near walking trails and large animal enclosures, we released all snakes at a standard location just outside of and uphill from the park (16°43’13.80" N, 93° 5’53.90" W). We collected and marked animals during 2004 and 2005 and continued recording recaptures via tattoos through 2009. When recapturing an animal, we proceeded as for initial captures, except we did not tattoo the animal again.

After mapping capture and recapture locations, we noticed clustered individual localities for 16 recaptures. To assess if the direction of recapture related to the previous capture, we calculated all 16 angles between a capture, the release point, and the next recapture. We then calculated 10,000 random distributions of 16 values distributed between 0–98 as determined by the maximum angle (98°) between the northwestern-most (16°43’25.24" N, 93° 5’59.03" W) and eastern-most (16°43’17.54" N, 93°5’39.06" W) points in the park from the release point (Fig. 1). We calculated the point probability of the observed data against the randomly generated data to measure what probability of this distribution fell beyond that value. With an alpha = 0.05, a value < 0.025 in a two-tailed distribution would reject the null hypothesis that observed data were sampled from the random distribution.

**Figure 1.**
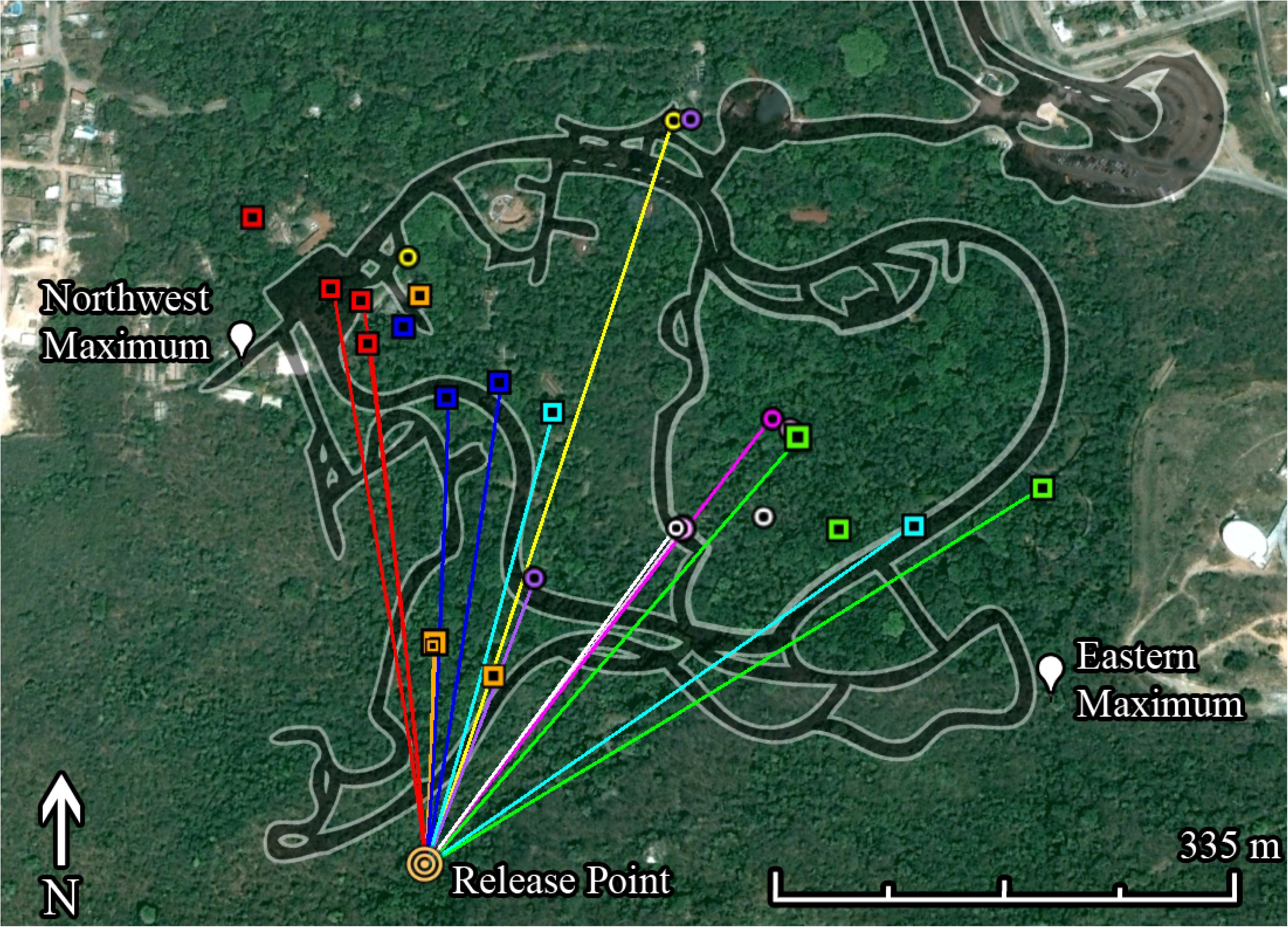
Adult *Micrurus apiatus* may maintain home ranges. Line and point colors correspond to individuals and minimum distances traveled to recapture locations. Points not associated with a line show initial captures. Square points are males and circular points are females. An outline of paths within the park is shown in black; however, to increase visibility it is not drawn to scale. The common release point for captured snakes and maximum boundaries used for random angle distribution generation are also shown.

For recaptures during all years, we calculated the minimum distance in meters traveled by animals from the common release point to the recapture point. This represents a minimum linear traveled distance and may vary from the actual distance traveled in this period of time (Pittman et al., 2014). We tested for several potential differences in minimum inferred movement between male and female snakes using Mann-Whitney U tests (Mann & Whitney, 1947; Table 1). Before distances or angles were calculated we converted all decimal degree coordinates to UTM (Synnatschke, 2016). We performed all statistics in R 3.1.3 (R Core Team, 2015). Data and R scripts were deposited in the Dryad repository: http://doi.org/XX.XXXX/X.

**Table 1.**
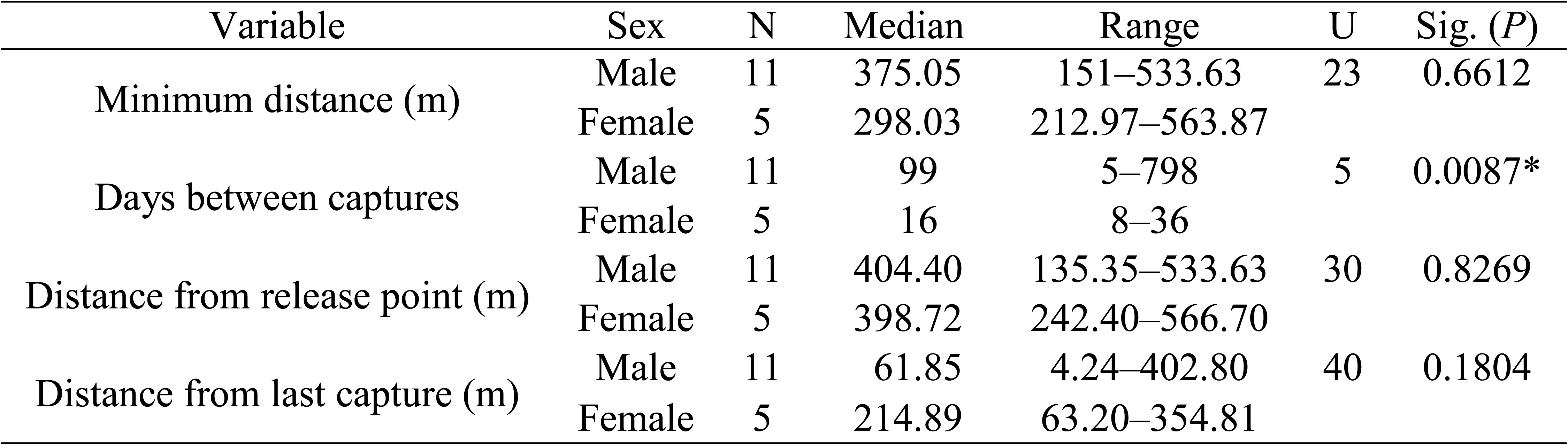
Recapture Mann-Whitney U comparisons for inferred traveling times and distances between sexes. * *P* < 0.05

## Results

In total, we observed 67 snakes, including 64 captures over six years of daily opportunistic sampling. Of the 64 captured snakes, we recaptured 10 individuals at least once within the park (Fig. 1). For recaptured animals, the capture-release-recapture angle (mean = 13.88587) was significantly less than the simulated dataset from 10,000 mean capture-release-recapture angles (mean = 49.03284, SD = 7.074059; point probability = 2.459777 × 10^−7^; Fig. 2). Observed data were 4.968 standard deviations less than the mean of simulated data (Fig. 2). Male (median = 9.957, N = 11) and female (median = 2.682, N = 5) snakes did not differ in capture-release-recapture angle (*U* = 20, *P* = 0.4409).

**Figure 2.**
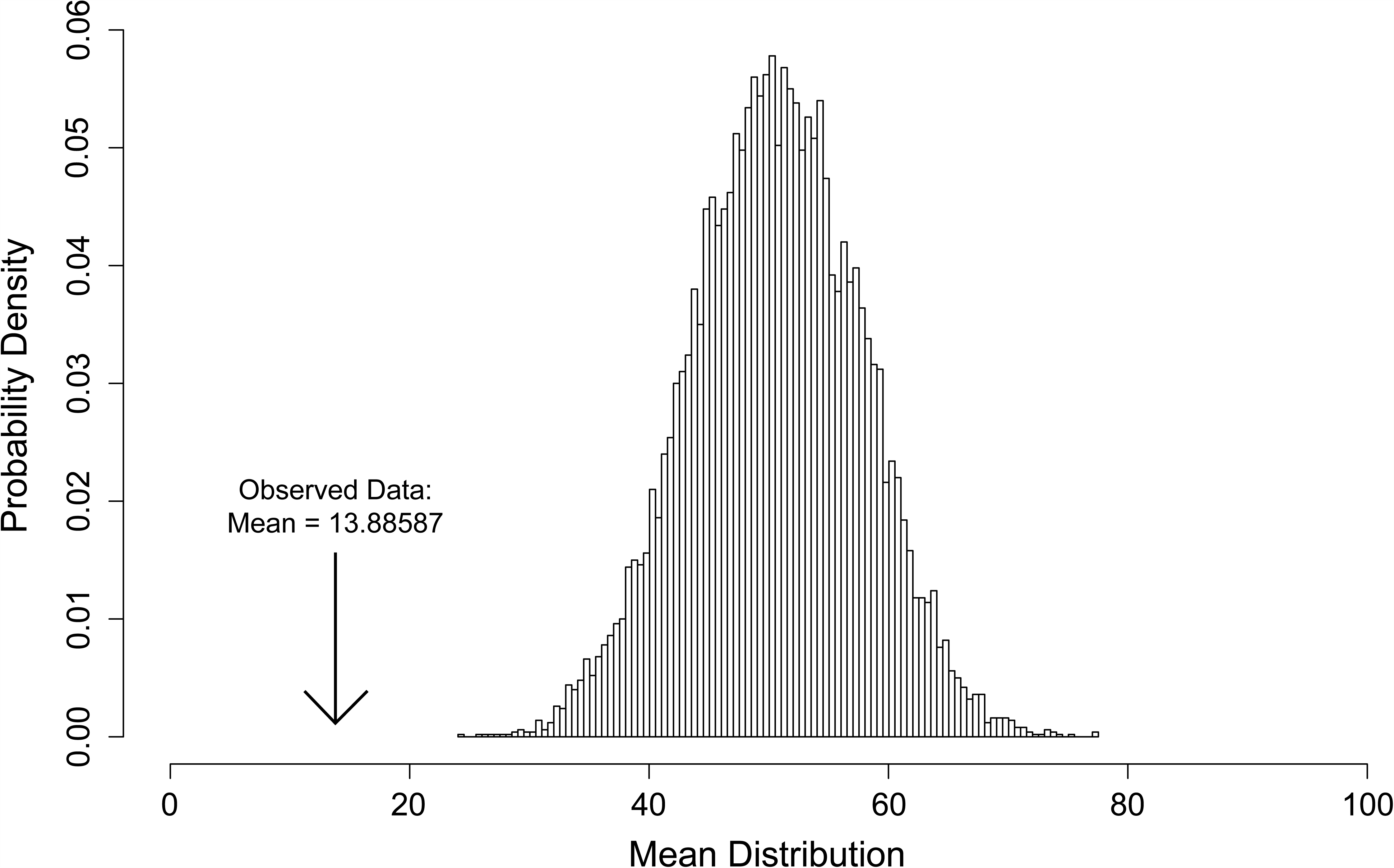
Probability distribution of 10,000 mean angles between *Micrurus apiatus’* capture point, release point, and last point of capture. Each angle distribution was randomly generated with a range of 0–98° and N=16 before calculating a mean. Data are shown binned by integers from 0–98. The mean of observed data is shown for comparison.

We recaptured males up to three times – all recaptured females were collected only a second time. Female snakes were recaptured sooner to their last capture than male snakes (Table 1). Neither the minimum distance traveled from a release point, distance captured from the release point, or the distance from last capture significantly differed between male and female *M. apiatus* recaptures (Table 1). Across all recaptured adult *M. apiatus*, snakes were recaptured closer to their original capture location with time (Fig. 3).

**Figure 3.**
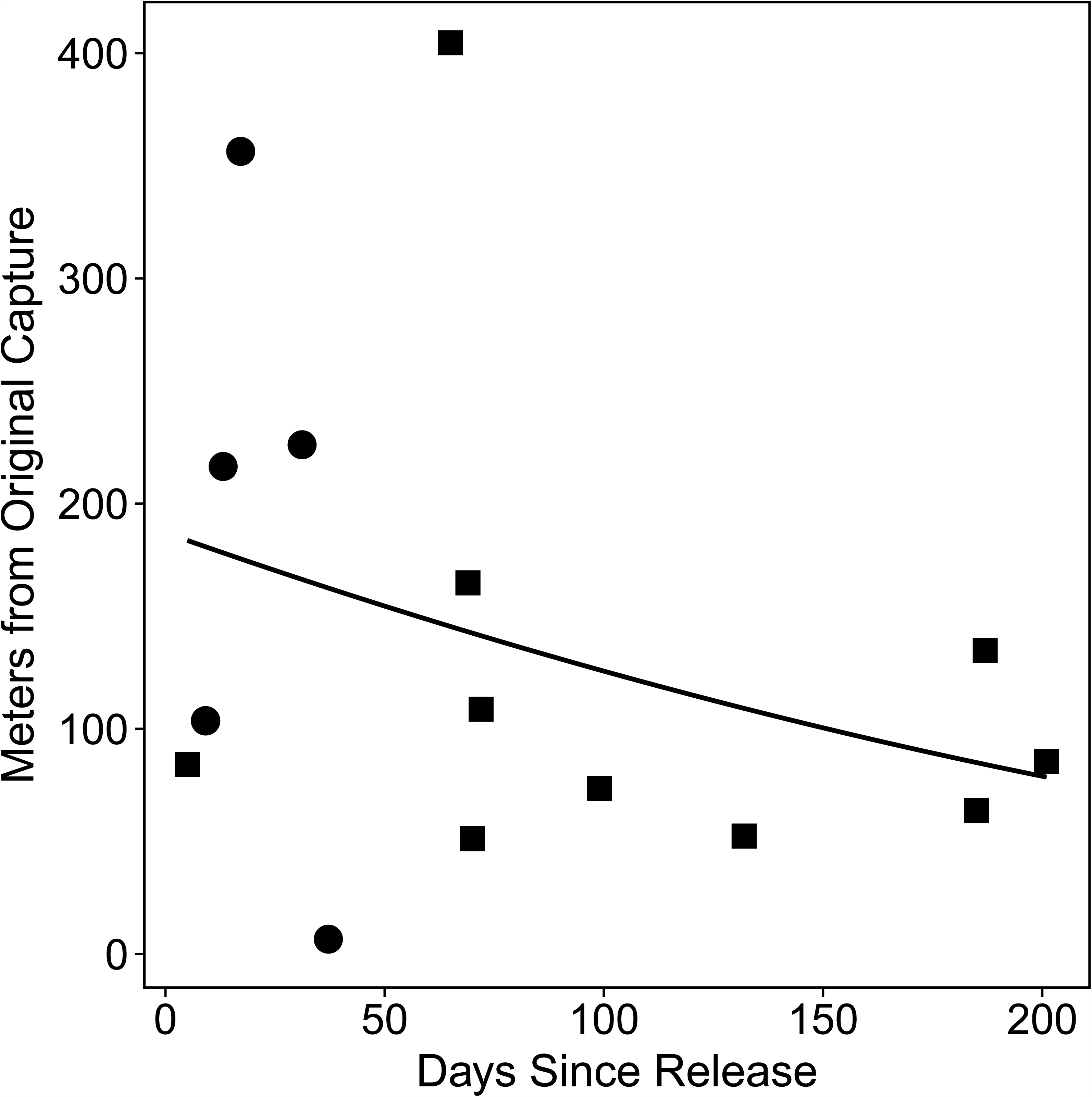
Generally, adult *Micrurus apiatus* returned to an original point of capture as time passed since their last release at a standard location. Square points are males and circular points are females. A best-fit line shows the polynomial formula: Meters = (0.0007398 ∙ Days^2^) – (0.6886 ∙ Days) + 183.03. One outlier was omitted: a male recaptured 798 days after previous capture.

## Discussion

For the first time, we report evidence of range maintenance and homing behavior in a coralsnake, an elapid from the new world (Figs 1–3). We captured *M. apatius* in a highly trafficked area. Since a *M. apiatus* bite can be lethal, we translocated animals to a common release point just outside of the park (Fig. 1). For this reason, we were surprised to find individual snakes close to their original capture points. In one example, we recaptured a male (red squares on Fig. 1) three times – all within an 75 m radius of the original capture point, but each at least 375 m from the release location. Furthermore, this animal’s second recapture was 419 meters from the release location after only five days. This observation implies a minimum dispersal capability of 83 meters per day, at least over less than a week.

We do not yet understand what distinguishes the microhabitats revisited by recaptured animals. Furthermore, we cannot present a mechanistic explanation for the high concordance between recapture locations. As snakes are highly reliant on olfactory cues for navigating microhabitats (Ford & Low, 1984; Mason, 1992; Rodda & Phillips, 1992), a chemosensory mechanism seems parsimonious. Amphibious sea kraits were found to home in open marine environments, suggesting that a chemical cue is unlikely to be the only mechanism involved in elapid philopatry (Shetty & Shine, 2002). Alternative explanations include a range of senses available to the snakes: auditory, tactile, visual, or possibly even magnetic (Rodda & Phillips, 1992; Southwood & Avens, 2010).

A recent study by Pittman et al. (2014) found an exceptionally similar pattern of putative “map sense” behavior in Burmese pythons (*Python molurus bivittatus*) introduced to the Florida everglades. These pythons likely used local cues to infer the direction of travel, though the authors leave open the possibility for a magnetic ‘compass sense.’ The paths from their capture locations to a common processing site and release site differed between animals, and each animal returned to near its original capture location for several days in succession, inferred from telemetry. Additionally, Hart et al. (2015) found seasonal home range maintenance in the same population of introduced Florida everglades pythons. Unfortunately, the mechanism for homing in pythons and snakes in general remains unknown. Considering *Micrurus* last shared a common ancestor with *Python* approximately 114-95 Myr (Noonan & Chippindale, 2006; Wiens, Brandley & Reeder, 2006; Vidal et al., 2009) it might be surprising if both taxa used the same mechanism. Homing has not yet been reported in other families of snakes, just in the Pythonidae and Elapidae. Further investigation of the distribution and qualities of homing behavior mechanisms in snakes may reveal enough data for a phylogenetic analysis of the loss and gain of shared mechanisms used for homing, if they are not all the same. Inferring patterns of evolution of snake homing underlying physiology will give us a firmer understanding of this behavior.

In a study apparently overlooked by Pittman et al. (2014), Singh et al. (1992) suggested the possibility of philopatry in an individual *Python molurus*. This python appeared to learn a home territory created by the authors and in one case returning after an absence of 9 days. Previous studies of terrestrial snake navigation after translocation suggest an absence of homing behavior. Instead, rattlesnakes (Reinert, 1991) and hognose snakes (Reinert & Rupert, 1999) in natural observation studies increase movement and likelihood to cross previously traveled paths, failing to demonstrate homing behavior.

In the present study, recaptured male and female *M. apiatus* moved about the same distance between captures, but from minimum distances traveled averaged across days, time between captures, and number of recaptures per animal we infer males move more aboveground than females (Table 1). It appeared that female *M. apiatus* were mostly found aboveground in a relatively short span of time compared to male snakes. Since the park was sampled daily, we do not think that this represents a sampling bias for increased probability of detection during certain times of the year due to methodology. A more complete analysis of the 67 captures will follow the present study, with additional comments on life history potentially generating this disparity between male and female recaptures. Data on the daily movements of elapids are few (Croak et al., 2013) or are restricted to one or two animals (e.g., Akani et al., 2005; Mohammadi et al., 2014), so it is difficult to compare against other studies in our assumption that snakes moved only the minimum linear distance between release and recapture. All data pertaining to coralsnake behavior and ecology can be considered valuable in consideration of widespread anthropogenic habitat destruction and global reptile declines.

## Acknowledgements

We thank the continuous efforts by ZOOMAT park staff in sighting these cryptic snakes. This project was conducted using best animal care practices of the Mexican state of Chiapas.

